# Rapid emergence of HCMV drug resistance in immunocompromised paediatric patients detected using target enrichment and deep sequencing

**DOI:** 10.1101/055814

**Authors:** Charlotte Houldcroft, Josephine M. Bryant, Daniel P. Depledge, Ben K. Margetts, Jacob Simmonds, Stephanos Nicolaou, Helena Tutill, Rachel Williams, Austen Worth, Stephen D. Marks, Paul Veys, Elizabeth Whittaker, Judith Breuer, the PATHSEEK condortium

## Abstract

**Background:** Cytomegalovirus can cause fatal disease in immunocompromised patients. With the advent of new anti-HCMV drugs there is interest in using virus sequence data to monitor resistance and identify new mutations.

**Methods:** We used target-enrichment to deep sequence HCMV DNA from 11 immunosuppressed paediatric patients receiving single or combination anti-HCMV treatment, serially sampled over 1-27 weeks. Changes in consensus sequence and resistance mutations were analysed for three ORFs targeted by anti-HCMV drugs and the frequencies of drug resistance mutations monitored.

**Results:** Targeted-enriched sequencing of clinical material detected mutations occurring at frequencies of 2%. Seven patients showed no evidence of drug resistance mutations. Four patients developed drug resistance mutations a mean of 16 weeks after starting treatment. In two patients, multiple resistance mutations accumulated at frequencies of 20% or less, including putative resistance mutations P522Q (UL54) and C480F (UL97). In one patient, resistance was detected 14 days earlier than by PCR. Phylogenetic analysis suggested recombination or superinfection in one patient.

**Conclusions:** Deep sequencing of HCMV enriched from clinical samples excluded resistance in 7 of eleven subjects and identified resistance mutations earlier than conventional PCR-based resistance testing in 2 patients. Detection of multiple low level resistance mutations was associated with poor outcome.

## Introduction

Cytomegalovirus (HCMV) is a ubiquitous betaherpesvirus with significant disease-causing potential in immunocompromised patients, including children with congenital immune deficiencies or immune suppression following solid organ or bone marrow transplantation. As well as causing pneumonitis, colitis, retinitis and uveitis[1] all of which contribute to HCMV-related mortality, HCMV disease also increases the risk of allograft vasculopathy and graft rejection, and significantly increases treatment costs[2]. Children are at particular risk from HCMV, with over 25% of primary HCMV infections in the UK occurring in childhood[3]. Up to 16% of patients on prolonged anti-HCMV therapy develop drug resistance[4, 5], many of them with mutations which cause multi-drug resistance [6]. However, it may be that not all the mutations that cause resistance are known and this may lead to underestimation of drug resistance in patients failing therapy.

Three drugs are currently licensed for HCMV prophylaxis and treatment, including ganciclovir, foscarnet, cidofovir; brincidofovir (the oral derivative of cidofovir) and letermovir are in phase III clinical trials; maribavir is available on a compassionate use basis. Treatment failure occurs in between 20[7]-50[8]% of HCMV cases, necessitating drug changes and in some cases the use of adoptive immunotherapy. Genetic evidence of drug resistance can guide clinical decision making[9] but current methods have technical limitations. Sanger sequencing of PCR amplicons only reliably detects drug resistance mutations that are present at frequencies of 20% or more[10]. Deep sequencing of PCR amplicons has enabled detection of minority resistance variants at frequencies as low as 1%[11] which could lead to earlier detection of HCMV resistance and better treatment. However, PCR and nested PCR are known to generate mutations which could make the identification of low level resistance mutations more difficult[12]. To minimize this problem, and to capture the genes currently implicated in antiviral resistance simultaneously, we made use of novel pulldown methodologies[12] and deep sequencing to analyse the UL27, 54 and 97 genes in serial samples from patients with prolonged HCMV viraemia despite anti-HCMV therapy. In this study, we include 11 retrospectively identified patients from Great Ormond Street Hospital for Children who had high CMV loads for two weeks or longer, with clinician suspicion of anti-viral drug resistance. By sequencing multiple samples from each patient we identified resistance mutations that were missed by conventional PCR and detected one mutation two weeks earlier. For the majority of patients, deep sequencing provided reassurance that antiviral resistance had not developed. Two patients who rapidly developed fixed resistance cleared virus following a change in treatment. However development of multiple sub-fixation resistance mutations in two patients was associated with poorer outcome.

## Methods

### Ethics and sample collection

Whole blood samples were stored at Great Ormond Street Hospital for Children (GOSH) at −80C. These residual samples were collected as part of the standard clinical care at GOSH, and subsequently approved for research use through the UCL Partners Infection DNA Bank by the NRES Committee London Fulham (REC reference: 12/L0/I089) All samples were anonymised. Eleven patients with CMV viral loads that remained unchanged or rose despite 2 weeks of first line anti-CMV therapy were selected. 20 samples from six patients (B [5], C [2], H [4], I [4], J [4] and M [1]) were tested for UL97 and/or UL54 resistance mutations by PCR and Sanger sequencing at the reference laboratory. Samples with sufficient material for DNA extraction (200μl) were analysed.

### DNA extraction, library construction, targeted enrichment, and sequencing

Total DNA was extracted from 200μl each sample using the EZ1 Virus kit and EZ1 XL extraction system (Qiagen) or DNA Blood Mini kit (Qiagen) according to manufacturer's instructions. Virus loads were established by an in-house NHS diagnostic qPCR assay (GOSH). To determine IU/ml, the copies/ml value is divided by 4.

### SureSelectXT Target Enrichment: RNA baits design

A library of 120-mer RNA baits spanning 115 GenBank HCMV whole and partial genome sequences were designed using the PATHSEEK consortium's in-house PERL script. Baits specificity was verified by BLASTn searches against the Human Genomic plus Transcript database. The custom-designed HCMV baits were uploaded to SureDesign and synthesised by Agilent Technologies.

### SureSelectXT Target Enrichment: Library preparation, hybridisation and enrichment

Total DNA from clinical samples was quantified using the Qubit dsDNA HS assay kit (Life Technologies, Q32854) and between 200-500ng of DNA was sheared for 150 seconds, using a Covaris E220 focused ultra-sonication system (PIP 175, duty factor 5, cycles per burst 200). End-repair, non-templated addition of 3’ poly A, adapter ligation, hybridisation, PCR (12 cycles pre-capture and 18 or 22 cycles post capture) and all post-reaction clean-up steps were performed according to the SureSelectXT Automated Target Enrichment for Illumina Paired-End Multiplexed Sequencing 200 ng protocol (version F.2) on the Bravo platform Workstation B (Agilent Technologies). All recommended quality control steps were performed on the 2200 TapeStation (Agilent Technologies). Samples were sequenced using the Illumina MiSeq platform. The presence of a subset of novel SNPs was confirmed by Sanger sequencing of PCR amplicons (GATC, Germany; Source Biosciences, UK; Manchester Medical Microbiology Partnership, UK; PHE, UK; and the Royal Free Hospital Virology Department, UK).

### Sequence assembly and variant analysis

Reads were trimmed to remove adapter sequences. Total reads were mapped to the HCMV reference sequence Merlin (RefSeq ID NC_006273) ORFs UL27, UL54 and UL97 using CLC Genomics Workbench 8.0.3 (Qiagen). Minority variants were called if: the base was sequenced at least five times; the variant was present in at least five reads (including two forward and two reverse reads); and it was present at a frequency of at least 2% (or 1% for bases sequenced over 1000 times). The read direction filter significance was 0.05 and the relative read direction filter significance was 0.01. Variants were identified using published lists of HCMV resistance mutations[13–17].

### Phylogenetic analysis

Sequences were aligned using ClustalW[18] and manually corrected in MEGA6 if necessary. Phylogenetic reconstructions were performed using MEGA6 maximum likelihood analysis (Tamura-Nei model, 1000 bootstraps, default settings)[19]. Sequences from the following HCMV genomes were used: NC_006273.2 (Merlin), KU317610.1 (AD169), JX512198.1 (Davis), AY223527.1 (Towne), GU937742.1 (Toledo), KJ872542.1 (PAV21), HQ380895.1 (JHC), KJ361971.1 (UKNEQAS1), KJ426589.1 (Han), KP745728.1 (BE/4/2010), KP745718.1 (CZ/1/2011).

## Results

The duration of HCMV positivity and treatment for each of the 11 patients is shown in table 1. Using SureSelect target enrichment we recovered sequence mapping to the UL27, 54, and 97 genes directly from all the clinical diagnostic samples in a single reaction without the need for virus isolation or PCR of overlapping genome fragments. A sample read mapping plot for each ORF is shown in supplementary figure 1. Details of mapping and coverage relative to virus genome copies/ml blood are shown in supplementary table 1 and supplementary figure 5. From deep sequencing results we were able to stratify patients into two groups: those with no evidence of developing resistance mutations despite receiving long term antiviral treatment (A, C, D, G, J, K, L); and those patients who developed known HCMV resistance mutations: either fixed (H and M) or at low level (B and I). We plotted viral load, drugs received and mutations over time for each of these patients (figure 1; supplementary figure 2).

**Table 1.**
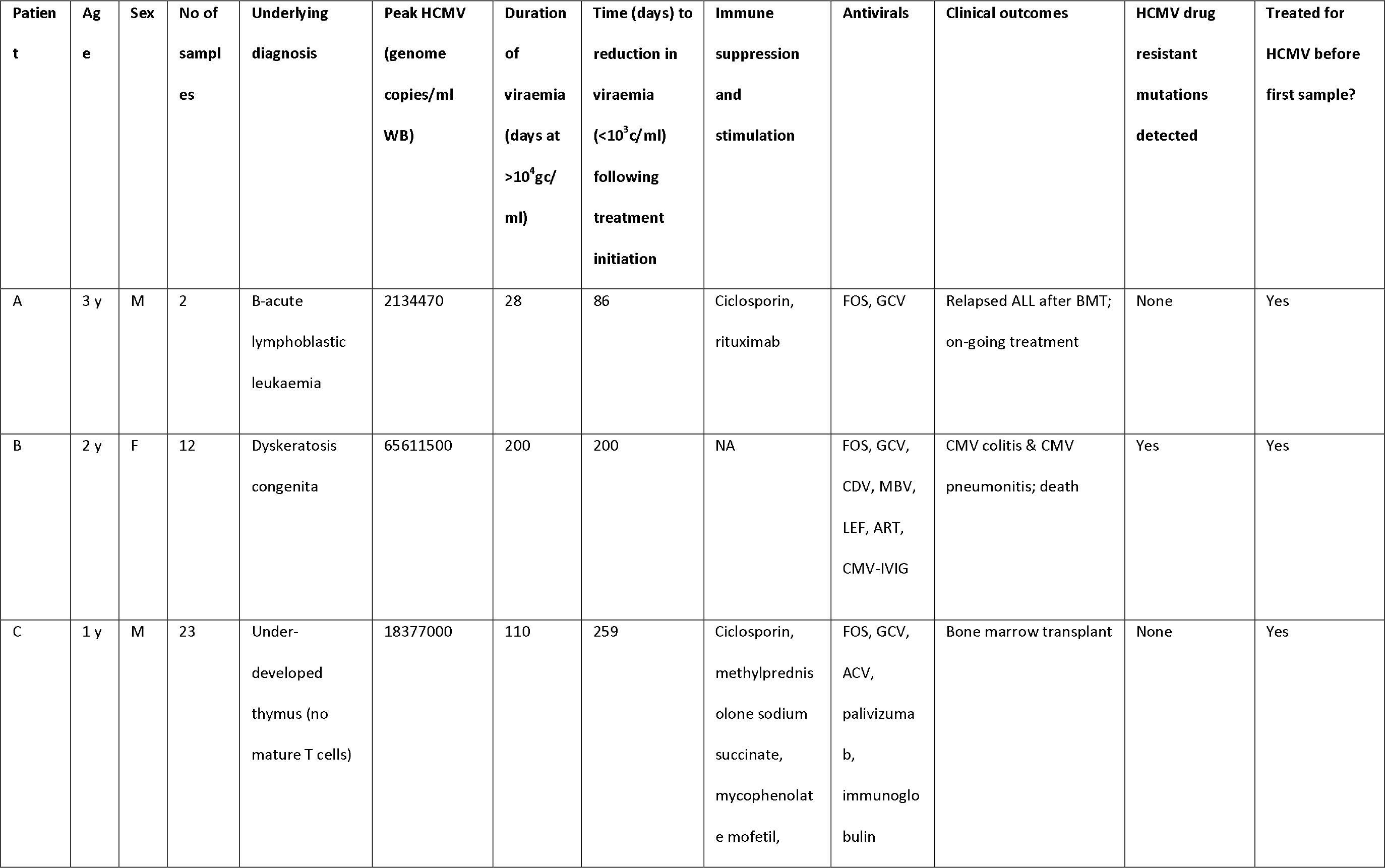
<u>Patient charecterstics</u>

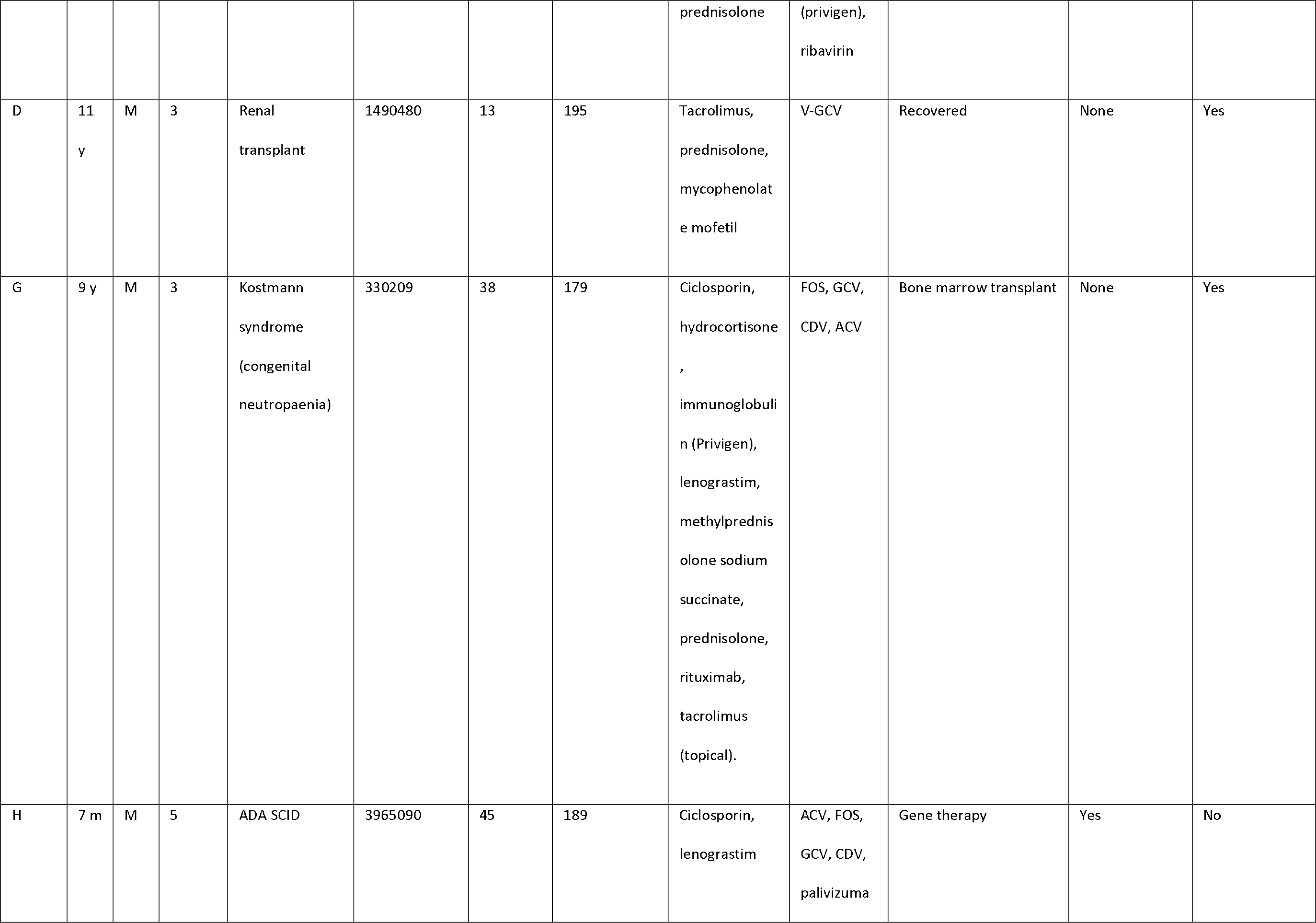

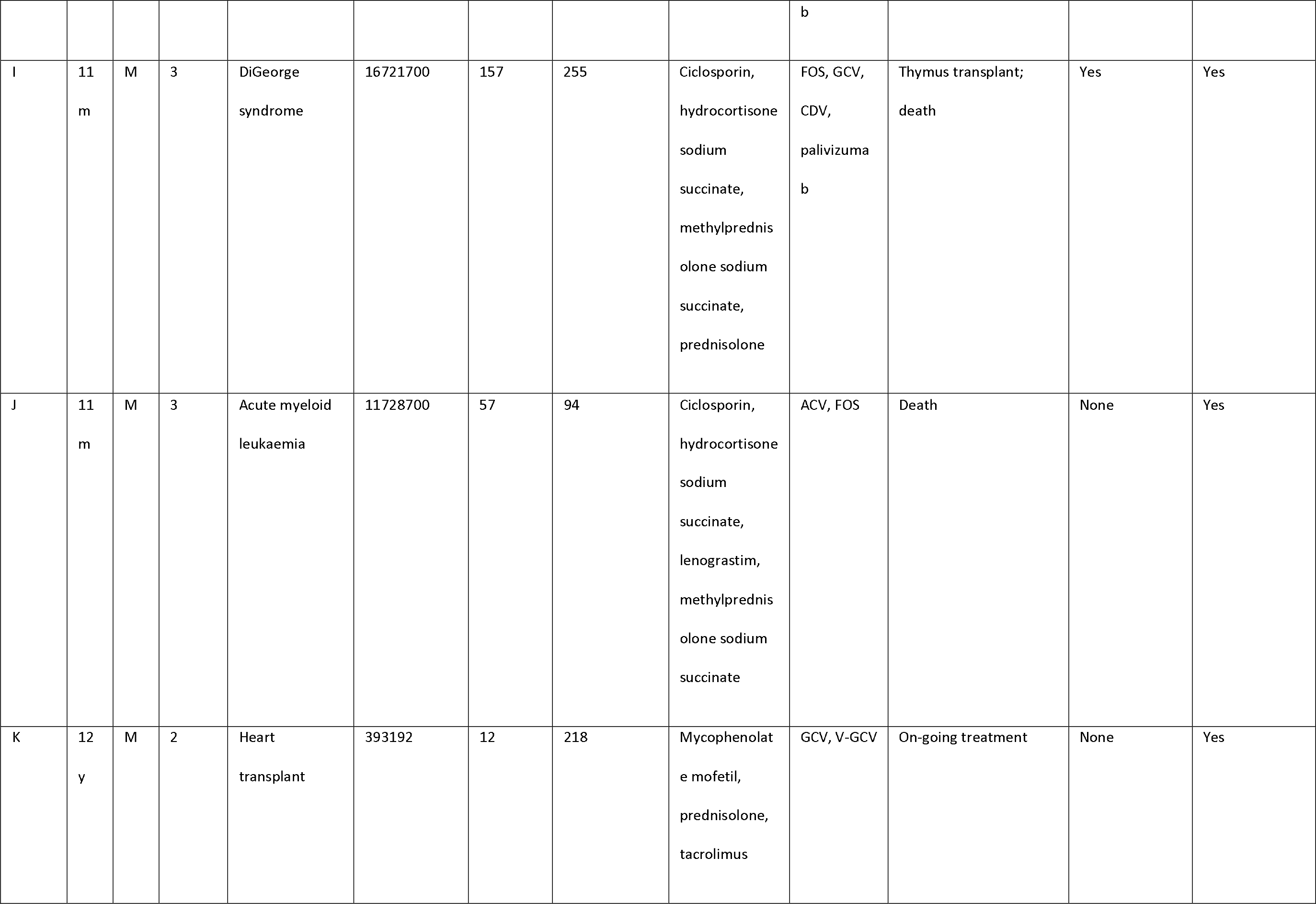

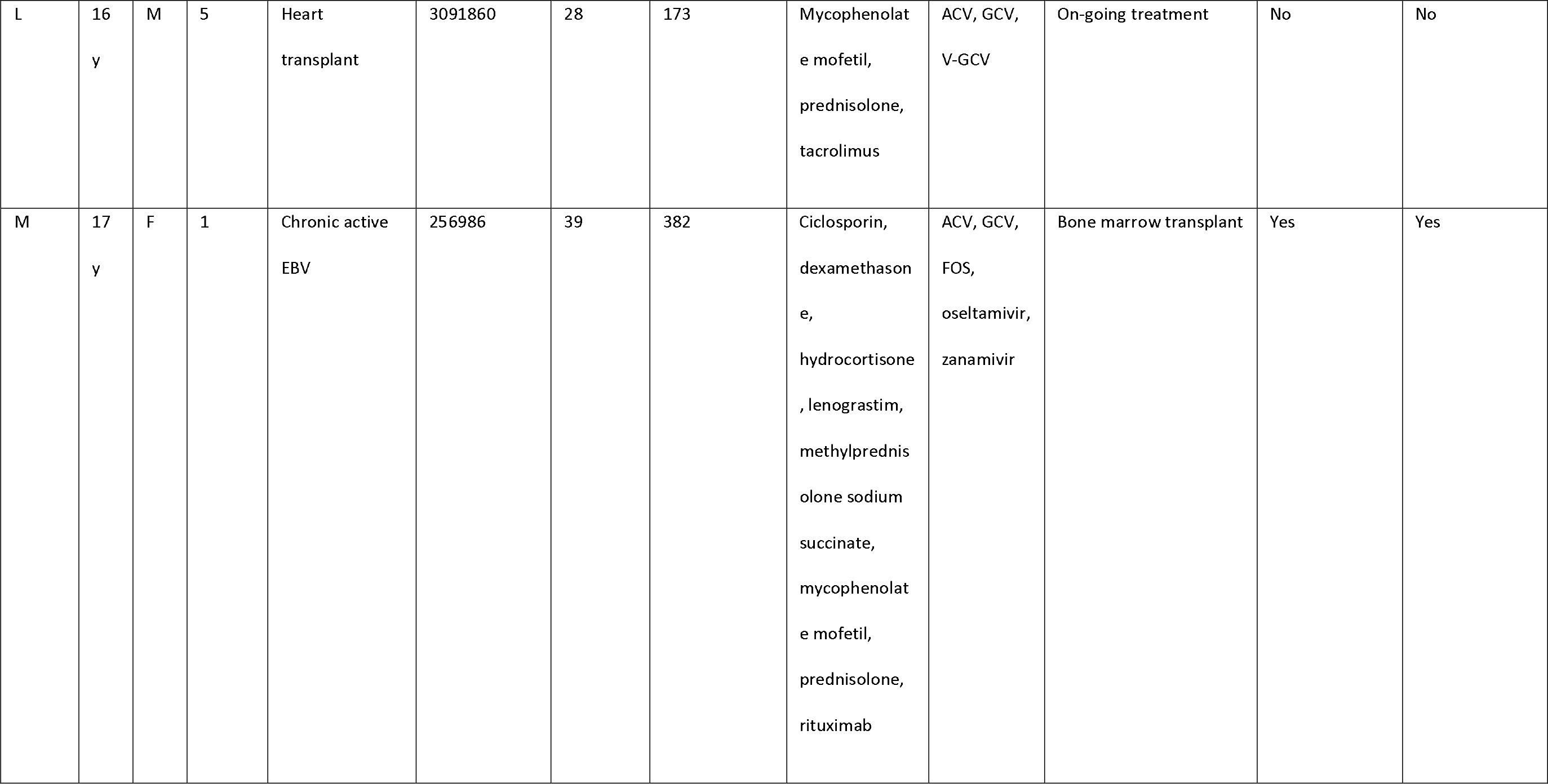

**Table 2.**
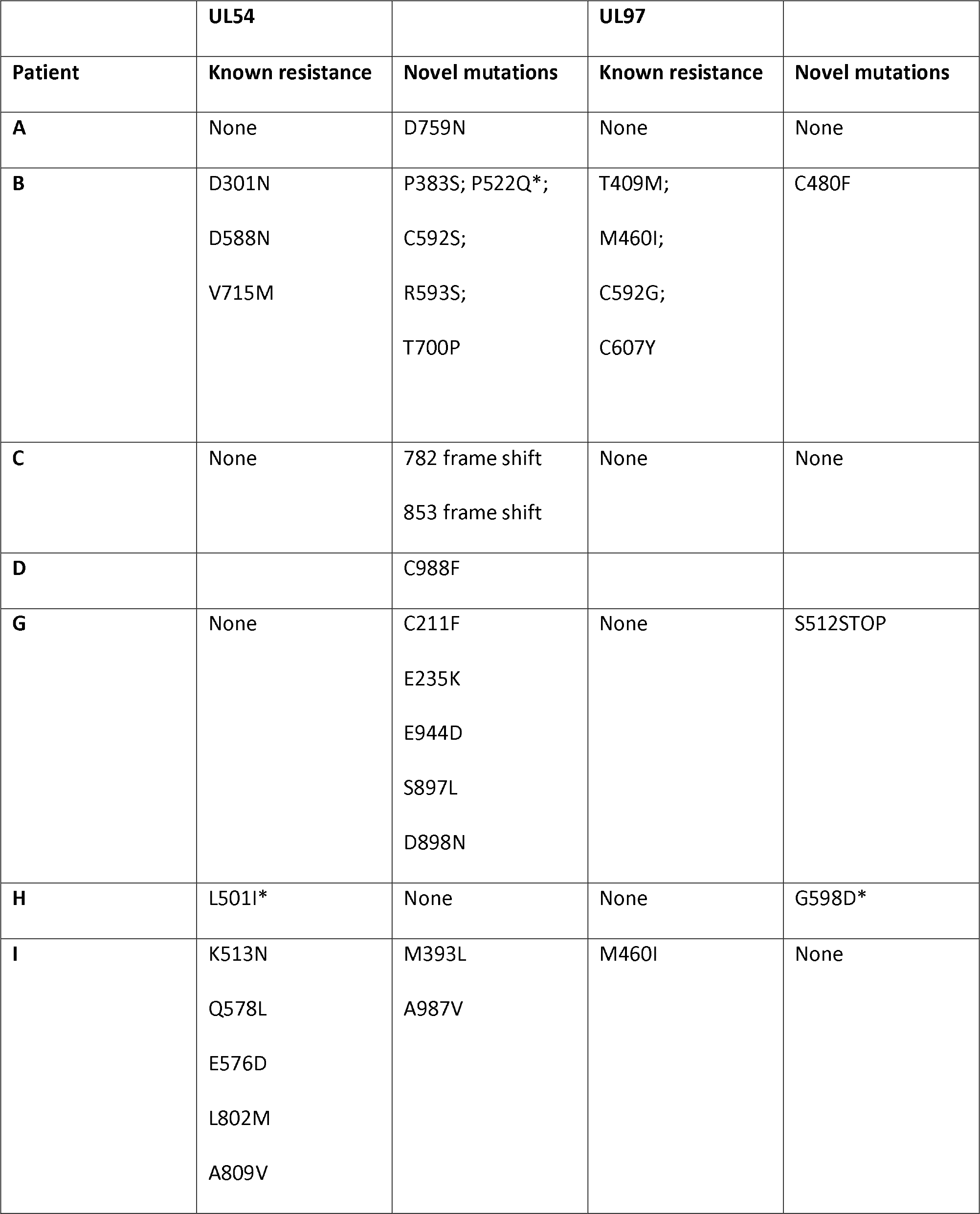
<u>Mutations</u>

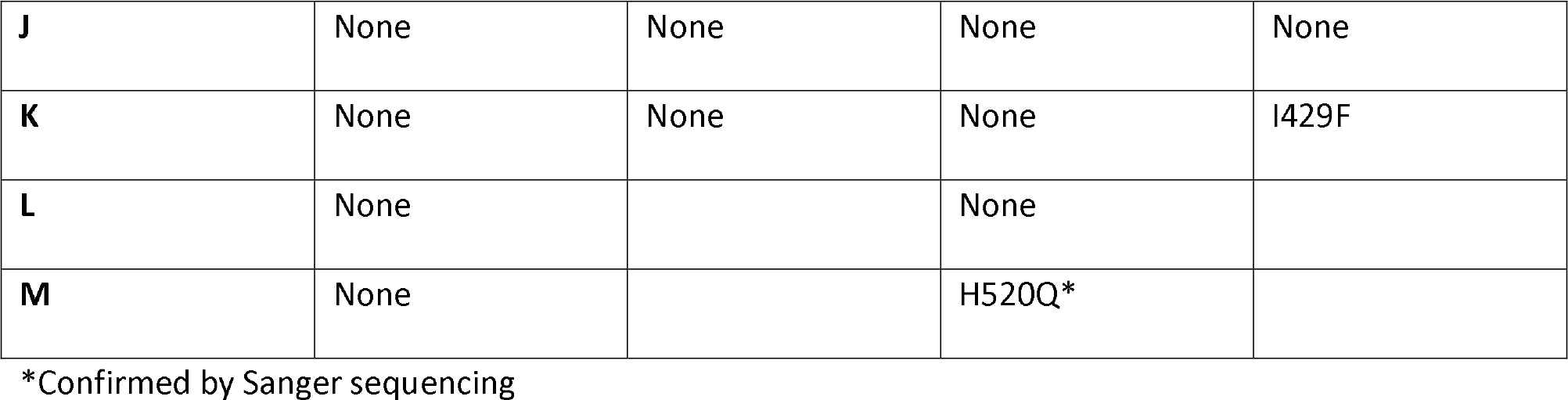

### Comparison of patients with and without drug resistance mutation

Comparing the four patients who developed resistance versus the seven who did not, the mean duration of treatment was longer in those who developed resistance (171 (SD 79) versus 101 (SD 70) days), the median number of antiviral drugs higher (3.5 versus 2), the peak viraemia higher (2.16×l0^7^ versus 5.36×10^6^ virus copies/ml blood) and mean duration of viraemia was greater (257 (SD 89) versus 172 (SD 63). Apart from the last (*p* = 0.048), the numbers were too small for these differences to be significant. Time to control of viremia in those who survived was faster in the two with resistance (118 (SD 47) versus 131 days (SD 85)). Mean total lymphocyte counts (TLC) were persistently low in patients B, I and J who died ie 0.46 (SD 0.56) as compared with patients A, D, G, H, K, L and M who survived and controlled their viremia to below 1000 copies/ml, mean TLC 1.26 (SD 0.85).

### Patterns of resistance mutations

Patients B, H, I and M developed known drug resistance mutations in UL54 and UL97 during treatment (figure 1). The mean time to mutations detection was 115 days (range 18-171) following the start of antiviral treatment. The mutations detected are shown in figure 2. Patient H carried no baseline resistance mutations by deep-sequencing analysis, but Sanger sequencing detected fixed resistance mutation L501I (CDV and GCV resistance) in ORF UL54 on day 18 of treatment (day 43 post-admission). This mutation was not detected by Sanger sequencing on day 56 of treatment despite continued GCV; the patient also received FOS throughout this period. The patient developed a mutation, G598D, in UL97 on day 81 post-admission (treatment day 56) which has previously been seen in patients failing GCV therapy, detected by Sanger sequencing. However the phenotype of this mutation without concurrent UL54 mutations has yet to be demonstrated by marker transfer[18, 19]. These samples were not available for follow-up deep-sequencing.

Patient M responded to FOS treatment with a reduction of viral load from ~250k copies/ml to ~50k copies/ml over 4 days, following the failure of GCV therapy caused by the fixed UL97 mutation H520Q known to cause an 8-fold (or greater) increase in GCV resistance[17, 20]. This mutation persisted at fixation for at least 43 days after GCV therapy was withdrawn.

In contrast, patients B and I developed multiple low frequency UL54 and UL97 drug resistance mutations after 112 and 171 days of treatment respectively. Neither patient was able to control their HCMV, and both eventually died of HCMV-related complications. In patient I, the first resistance mutation at position A809V in UL54 which is associated with HCMV growth rate attenuation[21] was detected at a frequency of 26% 171 days after starting GCV. This mutation was lost (or was present at a frequency of less than 5%) following withdrawal of GCV. The cessation of GCV and start of FOS and CDV was accompanied by a rise in the GCV UL97 resistance mutation M460I to 96% together with the UL54 resistance mutations Q578L (~3 fold FOS resistance) and K513N (12-fold CDV resistance) which rose to over 80% within 46 days. This pattern suggests that the M460I GCV resistance mutation was linked on the same virus to the UL54 resistance mutations which were selected for by FOS and CDV. The rising frequency of UL54 mutations was accompanied by a rise in HCMV load from 10^4^ to 10^7^ gc/ml. Patient I died with evidence of extensively drug-resistant HCMV, carrying multiple fixed and low frequency resistance mutations to CDV, GCV and FOS. PCR based resistance testing did not detect resistance until day 225, when only K513N was detected; on day 238, L802M was also detected as a ‘mixture’ by PCR.

A similar picture emerged in Patient B. Although resistance mutations were not detectable at >2% until after day 84 following the start of treatment, multiple low frequency (<40%) resistance mutations to GCV, FOS and CDV, with which the patient had been treated rapidly developed thereafter (figure 1). PCR and Sanger sequencing failed to detect these low level resistance mutations, with the exception of the GCV (D588N) substitution which was picked up 14 days after it became detectable by target enrichment. Despite a persistently high and increasing viral load, none of the low level resistance mutations rose to fixation (peak frequency <45%). The introduction of MBV, resulted in decline of the majority of low frequency GCV, FOS and CDV mutations (D301N, D588N and V715M in UL54 and M460I, C592G and C607Y in UL97). In contrast T409M in UL97 rose in frequency from 2% on day 175 (43 days after commencing MBV) to 39% at the point of treatment withdrawal. Mutation T409M is known to confer cross-resistance to MBV and GCV.

### Putative novel drug resistance mutations

Potential new resistance mutations were only seen in patient B. Mutations P522Q in UL54 and C480F in UL97 were detected at days 119 (P522Q) and 175 (C480F), ie 14 days before and 42 days following the introduction of maribavir, respectively, with the former increasing to 84% by day 193 (60 days following the start of maribavir treatment) and the latter also increasing over time. P522Q and C480F have not previously been reported as resistance mutations although variants P522S and P522A are associated with GCV and CDV resistance[22], and C480R is associated with increased resistance to methylenecyclopropane nucleoside analogues[23]. C480F appeared at a frequency of 5% at approximately the same time as the known MBV mutation T409M, rising to 58% by day 193 (60 days of MBV treatment). P522Q appeared first of the previously undetected mutations and rose rapidly to fixation following initiation of MBV treatment. The appearance of these three mutations was accompanied by rising viral load, suggesting that all three may confer resistance to MBV.

## Stop codons, insertions and deletions

Patients I and L (despite never having received MBV) showed evidence of fixed truncating mutations in UL27 (supplementary figure 4i) both of which would be predicted to confer resistance and/or growth attenuation [13, 24], In patient G, a minority stop codon (~10%) was detected at amino acid position 512 in UL54 day 63 post-admission, but was not detected in subsequent samples from this patient (supplementary figure 4ii).

In samples from a number of patients, we detected low-frequency frame shift mutations in ORF UL54, at frequencies of between 2 and 13%: A (<5%); B (<10%); C (<6%); D (<12%); H (10%); and K (13%) (supplementary figure 3). Many of these mutations were lost over time, or replaced by different frame shifts, suggesting they are unfit.

### Phylogenetic analysis of sequences from patients with multi-drug resistance

To examine further the complex drug resistance patterns seen in patients B and I, we constructed a phylogenetic tree for each of the three target regions, including all samples from these patients and eleven publically available HCMV genomes from GenBank (figure 3i-iii). For patient B, UL27 consensus sequences clustered in different parts of the tree in a time dependent manner (figure 3i). The consensus sequences of genes UL54 and UL97 show change over time in patients B and I that is compatible with sequence evolution due to anti-viral drug pressure. In Patient B the changes in phylogenetic clustering for UL27 occurred after the start of maribavir on day 133, and may reflect recombination or re-infection with a second strain of HCMV in this patient.

## Discussion

Persistent HCMV viraemia is associated with poor outcomes in immunosuppressed patients, including those undergoing bone marrow[2] and solid organ transplantation, and treatment with anti-HCMV drugs is indicated. HCMV viraemia carries significant economic costs, estimated at £22,500 ($32000) per paediatric bone marrow transplant patient[2]. To explore treatment failure, testing for resistance mutations and if necessary a change in therapy is recommended if the viral load remains the same or rises after two[25] or three[8] weeks of treatment. Changes in treatment may also be prompted by side effects, and bone marrow function particularly in haematological stem cell transplant recipients. In this study we used deep sequencing to investigate drug resistance patterns in persistently viraemic patients requiring prolonged treatment. Notwithstanding persistent viraemia, seven patients showed no sign of drug resistance and six of them were able to control their viremia to below 10^3^ gc/ml while on treatment. Patients who developed resistance had higher viremia, lower lymphocyte counts more drugs and longer duration of antiviral treatment, although numbers were too small for these differences to be significant. Overall, these data support the findings of others, that development of drug resistance mutations are associated with poor control of viremia and represent a poor prognostic indicator in immunosuppressed patients receiving treatment for HCMV; two of four patients developing resistance mutations died as compared with one of seven who remained resistance-free. Notwithstanding these findings, the two patients H and M, in whom resistance mutations rose rapidly to fixation, responded to a change in treatment and controlled their viremia (two qPCR results <10^3^ gc/ml) within a mean of 17 weeks. In patient M the H520Q resistance mutation to GCV in ORF UL97 persisted despite withdrawal of the drug, suggesting that this variant remained fit despite the H520Q mutation.

By contrast, where we identified multiple mutations occurring simultaneously, in patients B and I, this was associated with profound treatment failure and death from HCMV-related disease. Observations from deep-sequencing of PCR amplicons suggest that multiple resistance mutations occurring at sub-fixation levels can contribute to a drug-resistant phenotype and this is consistent with the evidence,
particularly in patient B in whom high HCMV viral loads persisted in the presence of multiple often low frequency mutations (figure 1). One explanation is that low frequency drug resistance mutations are distributed throughout the viral population resulting in many relatively unfit resistant viruses, none of which can outcompete the others[26]. A similar pattern in seen in patient I, in whom a change from GCV to CDV appears to have selected for one set of resistance mutations in favour of another, perhaps because these mutations arose on different populations of the virus within this patient. Further evidence for this comes from mouse studies making use of cells infected with multiple murine CMV strains. These strains trans-complement one another, increasing overall viral fitness[27]. Both patients I and B showed a rapid rise in resistance mutations in response to treatment changes, with concomitant loss of others. This pattern, particularly in patient B for whom more samples were available, is consistent with low level persistence of multiply resistant viruses which rapidly replicate under the selective pressure of a new drug. Conventional PCR and Sanger sequencing failed on at least two occasions to detect any of these mutations, apart from the D588N which was present at a frequency of 9%. Deep sequencing is therefore able to detect potentially important drug resistance that is missed by conventional methods. For example the P552Q mutation was detected at frequencies of 1.67% (day 119), 4% (day 126) and 10.64% (day 133), prior to the start of maribavir on day 133. Similarly, PCR and Sanger sequencing of samples from patient I missed multiple drug resistance mutations at frequencies of 2-41%, 54 days after these mutations became detectable by target-enriched sequencing.

The speed with which the virus became resistant in patient B and the loss of four drugs resistant mutations in UL97 and UL54, suggested strain replacement rather *die now* mutation and prompted us to examine the possibility of mixed infection. This change in phylogenetic clustering for UL27 sequences following the introduction of marabivir confirmed this suspicion. HCMV is known to be highly recombigenic[35], and in this case, without whole genome analysis, we are unable to distinguish the possibility of recombination, re-infection, or reactivation of a pre-existing secondary strain. In summary we have demonstrated that deep sequencing of HCMV ORFs UL27, 54 and 97 could be achieved directly from whole blood with virus loads in the range 80,000-65,000,000 copies/ml without prior culture or PCR. We were able to detect resistance mutations occurring at 2% or more in patients with viraemia persisting at levels of ≥10^4^ gc/ml for two weeks or more. Our data suggest that in contrast to amplicon and Sanger sequencing, deep sequencing can exclude resistance in patients with persistently high levels of viraemia, thereby providing a measure of support prolonging current antiviral treatment or returning to them at a later date if further treatment is needed. Where resistance mutations are detected, we observed two patterns, rapid development of fixed resistance with clearance of virus following a change in treatment, and development of multiple sub-fixation resistance mutations, with potentially poorer outcome. Further investigation is needed to determine whether these patterns are indeed predictive of outcome. We do not yet understand why multiple minority drug resistance mutations arise in some patients. Multiple minority variants, which are likely to be better detected using deep sequencing methods, appeared to complicate treatment to a greater extent than single fixed resistance mutations. In our patients multiple low level drug mutations was associated with poor prognosis probably because they increased the risk that a change in drug would select for a preexisting mutation. Deep-sequencing of HCMV allows us to characterise these mutations and could be used to inform which drugs are given earlier in treatment, or to highlight those patients for whom additional non-pharmacological interventions such as withdrawal of immunosuppression, or the use of virus-specific cytotoxic T lymphocytes are most appropriate.

## Data availability

Raw sequencing data has been deposited in the European Nucleotide Archive under project accession PRJEB12814. Bait sequences are available by request from the authors.

## Author contributions

Judith Breuer (JB), CJH and DPD conceived the study design. CJH, EW, JS, AW, SM and PV supplied patient clinical data. CJH, DPD, and SN performed the DNA extractions. DPD, HT, CJH and SN sequenced the samples. RW administered the study. CJH, JMB and DPD analysed the data. CJH, JMB and JB wrote the paper. All authors read and approved the manuscript.

## Acknowledgements

We acknowledge all partners within the PATHSEEK consortium (University College London, Erasmus MC, QIAGEN AAR, and Oxford Gene Technology). We acknowledge infrastructure support from the UCL MRC Centre for Molecular Medical Virology. The clinical HCMV samples were provided by the UCL Infection DNA Bank, supported by University College London, Great Ormond Street Hospital, Royal Free Hospital and Barts and The London NHS Trust. This project was supported by the National Institute for Health Research Biomedical Research Centre at Great Ormond Street Hospital for Children NHS Foundation Trust and University College London. The authors would also like to thank Dr James Taylor (University of Cambridge), Dr Kimberly Gilmour (Great Ormond Street Hospital) and Dr Karen Buckland (UCL Institute of Child Health).

## Previous presentations

Data associated with this paper were previously presented at the Society for General Microbiology conference in Birmingham, UK (2015).

## Abbreviations

ART: Artusenate
BCDV: Brincidofovir
CAEBV: Chronic active EBV
CDV: Cidofovir
HCMV: Human cytomegalovirus
CMV-IVIG: Cytomegalovirus intravenous immune globulin
CTL: Cytotoxic T cells
FOS: Foscarnet
GCV: Ganciclovir
GOSH: Great Ormond Street Hospital for Children, UK
LEF: Leflunomide
LTV: Letermovir
MBV: Maribavir
VGCV: Valganciclovir
WB: Whole blood

## References

1. Rafailidis PI, Mourtzoukou EG, Varbobitis IC, Falagas ME. Severe cytomegalovirus infection in apparently immunocompetent patients: a systematic review. Virology journal 2008; 5:47.

2. Hiwarkar P, Gaspar HB, Gilmour K, et al. Impact of viral reactivations in the era of pre-emptive antiviral drug therapy following allogeneic haematopoietic SCT in paediatric recipients. Bone Marrow Transplant 2013; 48:803–8.

3. Patrick EJ, Higgins CD, Crawford DH, McAulay KA. A cohort study in university students: investigation of risk factors for cytomegalovirus infection. Epidemiol Infect 2014; 142:1990–5.

4. Shmueli E, Or R, Shapira MY, et al. High rate of cytomegalovirus drug resistance among patients receiving preemptive antiviral treatment after haploidentical stem cell transplantation. The Journal of infectious diseases 2014; 209:557–61.

5. Couzi L, Helou S, Bachelet T, et al. High incidence of anticytomegalovirus drug resistance among D+R-kidney transplant recipients receiving preemptive therapy. American journal of transplantation: official journal of the American Society of Transplantation and the American Society of Transplant Surgeons 2012; 12:202–9.

6. Hantz S, Garnier-Geoffroy F, Mazeron MC, et al. Drug-resistant cytomegalovirus in transplant recipients: a French cohort study. The Journal of antimicrobial chemotherapy 2010; 65:2628–40.

7. Asberg A, Humar A, Rollag H, et al. Oral valganciclovir is noninferior to intravenous ganciclovir for the treatment of cytomegalovirus disease in solid organ transplant recipients. American journal of transplantation: official journal of the American Society of Transplantation and the American Society of Transplant Surgeons 2007; 7:2106–13.

8. van der Beek MT, Marijt EW, Vossen AC, et al. Failure of pre-emptive treatment of cytomegalovirus infections and antiviral resistance in stem cell transplant recipients. Antiviral therapy 2012; 17:45–51.

9. Houldcroft C. Sequencing drug-resistant cytomegalovirus in pediatric patients: toward personalized medicine. Future Virology 2015:1–4.

10. Sahoo MK, Lefterova MI, Yamamoto F, et al. Detection of cytomegalovirus drug resistance mutations by next-generation sequencing. Journal of clinical microbiology 2013; 51:3700–10.

11. Gorzer I, Guelly C, Trajanoski S, Puchhammer-Stockl E. Deep sequencing reveals highly complex dynamics of human cytomegalovirus genotypes in transplant patients over time. J Virol 2010; 84:7195-203.

12. Depledge DP, Palser AL, Watson SJ, et al. Specific capture and whole-genome sequencing of viruses from clinical samples. PLoS One 2011; 6:e27805.

13. Hakki M, Drummond C, Houser B, Marousek G, Chou SW. Resistance to maribavir is associated with the exclusion of pUL27 from nucleoli during human cytomegalovirus infection. Antivir Res 2011; 92:313-8.

14. Chou S. Phenotypic diversity of cytomegalovirus DNA polymerase gene variants observed after antiviral therapy. J Clin Virol 2011; 50:287–91.

15. Chou S. Rapid In Vitro Evolution of Human Cytomegalovirus UL56 Mutations That Confer Letermovir Resistance. Antimicrob Agents Chemother 2015; 59:6588–93.

16. Chou S. Approach to drug-resistant cytomegalovirus in transplant recipients. Curr Opin Infect Dis 2015; 28:293–9.

17. Gohring K, Hamprecht K, Jahn G. Antiviral Drug-and Multidrug Resistance in Cytomegalovirus Infected SCT Patients. Comput Struct Biotechnol J 2015; 13:153–9.

18. Thompson JD, Gibson TJ, Higgins DG. Multiple sequence alignment using ClustalW and ClustalX. Curr Protoc Bioinformatics 2002; Chapter 2:Unit 23.

19. Tamura K, Stecher G, Peterson D, Filipski A, Kumar S. MEGA6: Molecular Evolutionary Genetics Analysis version 6.0. Molecular biology and evolution 2013; 30:2725–9.

20. Chou S, Waldemer RH, Senters AE, et al. Cytomegalovirus UL97 phosphotransferase mutations that affect susceptibility to ganciclovir. The Journal of infectious diseases 2002; 185:162–9.

21. Chou S, Marousek GI, Van Wechel LC, Li S, Weinberg A. Growth and drug resistance phenotypes resulting from cytomegalovirus DNA polymerase region III mutations observed in clinical specimens. Antimicrob Agents Chemother 2007; 51:4160–2.

22. Chou S. Cytomegalovirus UL97 mutations in the era of ganciclovir and maribavir. Reviews in medical virology 2008; 18:233–46.

23. Komazin-Meredith G, Chou S, Prichard MN, et al. Human cytomegalovirus UL97 kinase is involved in the mechanism of action of methylenecyclopropane analogs with 6-ether and-thioether substitutions. Antimicrob Agents Chemother 2014; 58:274–8.

24. Chou S. Diverse cytomegalovirus UL27 mutations adapt to loss of viral UL97 kinase activity under maribavir. Antimicrob Agents Chemother 2009; 53:81–5.

25. Boeckh M, Ljungman P. How we treat cytomegalovirus in hematopoietic cell transplant recipients. Blood 2009; 113:5711–9.

26. Chou S, Ercolani RJ, Sahoo MK, Lefterova MI, Strasfeld LM, Pinsky BA. Improved detection of emerging drug-resistant mutant cytomegalovirus subpopulations by deep sequencing. Antimicrob Agents Chemother 2014; 58:4697–702.

27. Cicin-Sain L, Podlech R, Messerle M, Reddehase MJ, Koszinowski UH. Frequent coinfection of cells explains functional in vivo complementation between cytomegalovirus variants in the multiply infected host. Journal of Virology 2005; 79:9492–502.

28. Drouot E, Piret J, Boivin G. Novel method based on “en passant” mutagenesis coupled with a gaussia luciferase reporter assay for studying the combined effects of human cytomegalovirus mutations. Journal of clinical microbiology 2013; 51:3216–24.

29. Manuel O, Asberg A, Pang X, et al. Impact of genetic polymorphisms in cytomegalovirus glycoprotein B on outcomes in solid-organ transplant recipients with cytomegalovirus disease. Clinical infectious diseases: an official publication of the Infectious Diseases Society of America 2009; 49:1160–6.

30. Coaquette A, Bourgeois A, Dirand C, Varin A, Chen W, Herbein G. Mixed cytomegalovirus glycoprotein B genotypes in immunocompromised patients. Clinical infectious diseases: an official publication of the Infectious Diseases Society of America 2004; 39:155–61.

31. Sowmya P, Madhavan HN. Analysis of mixed infections by multiple genotypes of human cytomegalovirus in immunocompromised patients. J Med Virol 2009; 81:861–9.

32. Stanton R, Westmoreland D, Fox JD, Davison AJ, Wilkinson GW. Stability of human cytomegalovirus genotypes in persistently infected renal transplant recipients. J Med Virol 2005; 75:42–6.

33. Puchhammer-Stockl E, Gorzer I, Zoufaly A, et al. Emergence of multiple cytomegalovirus strains in blood and lung of lung transplant recipients. Transplantation 2006; 81:187–94.

34. Puchhammer-Stockl E, Gorzer I. Human cytomegalovirus: an enormous variety of strains and their possible clinical significance in the human host. Future Virology 2011; 6:259–71.

35. Sijmons S, Thys K, Mbong Ngwese M, et al. High-throughput analysis of human cytomegalovirus genome diversity highlights the widespread occurrence of gene-disrupting mutations and pervasive recombination. Journal of virology 2015.

